# Wildfire affects circulating hormone levels and the expression of male sexual plumage in a tropical songbird

**DOI:** 10.1101/2020.12.13.422576

**Authors:** Jordan Boersma, Douglas G. Barron, Daniel T. Baldassarre, Michael S. Webster, Hubert Schwabl

**Author notes:** **Correspondence:** Jordan Boersma, School of Biological Sciences, Washington State University, Pullman, WA, 99164, USA.

## Abstract

Natural disturbances like drought and wildfires are expected to increase in prevalence, so understanding how organisms are affected is a key goal for conservationists and biologists alike. While many studies have illustrated long-term effects of perturbations on survival and reproduction, little is known of short-term effects to physiology and sexual signal expression. Ornamental traits have been proposed as reliable indicators of environmental health, yet studies are lacking in the context of natural disturbances. Here we present short-term responses of male Red-backed Fairywrens (*Malurus melanocephalus*) to wildfire near the onset of the typical breeding season. Males of this species are characterized by plastic expression of sexual plumage phenotypes in their first breeding season. Using two populations with Fairywren captures before and after separate wildfires we illustrate that wildfire suppressed molt into ornamented plumage, including in third year males that typically show little plasticity in ornamentation. Baseline plasma corticosterone was elevated in males sampled after fire, but condition (furcular fat stores) was unaffected. Although testosterone levels did not decrease following fire, we found a positive correlation between testosterone and plumage ornamentation. In addition, males molting in ornamental plumage had higher circulating levels of testosterone than males molting in unornamented plumage following fire. Collectively, these findings suggest that wildfires inhibit or greatly delay acquisition of ornamentation in young males without exerting obvious effects on condition, but rather through subtle effects on testosterone and corticosterone circulation. This natural experiment also reveals that expression of alternative male reproductive phenotypes in this species is sensitive to environmental conditions and more plastic than previously assumed.

## INTRODUCTION

As climate change increases the prevalence of drought (Dai 2013, Cook et al. 2014, Trenberth et al. 2014), wildfires are expected to increase in frequency and severity (Flannigan et al. 2013). Understanding the myriad effects wildfires produce for native fauna is thus of utmost importance. Many factors including seasonal timing and interval of fires, as well as whether species evolved in habitats that historically burn, dictate how organisms are affected by wildfire (Hobson and Schieck 1999, Woinarski 1999, Cox and Widener 2008, Stojanovic et al. 2016). Tropical birds are especially vulnerable to the threat of climate change (reviewed in Şekercioĝlu et al. 2012), yet few studies have explored how wildfires affect tropical species outside of Amazonian forests (Barlow et al. 2002, 2006, Barlow and Peres 2004a, b, Mestre et al. 2013, Lemos Da Silva et al. 2015). Determining how tropical species respond to environmental disturbances can establish which components of phenology are fixed and which are plastic, which has impacts both on conservation and life history evolution. For instance, a recent analysis of drought impacts on breeding bird communities in the old and new world tropics revealed that shorter-lived tropical species suffer from decreased survival and continue to breed while longer-lived tropical species delay breeding and thus buffer against detrimental effects to survival (Martin and Mouton 2020). Illuminating the mechanisms underlying the tradeoff between reproduction and survival is thus integral to understanding how organisms respond to natural disturbances such as wildfires.

Testosterone mediates the tradeoff between reproduction and survival in many bird species as it is essential for spermatogenesis and expression of mating behavior, but elevation of testosterone can come with a host of detrimental effects to survival via endogenous costs including immunosuppression and external costs like increased predation and injury (Cawthorn et al. 1998, Peters 2000, Wingfield et al. 2001, Roberts et al. 2004, Reed et al. 2006, McGlothlin et al. 2010, Foo et al. 2017). The expression of secondary sexual characters like plumage ornamentation are stimulated by testosterone in males of many bird species (reviewed in Kimball and Ligon 1999; Hau 2007), which can produce their own costs to survival. Sexual ornaments are often considered honest signals due predominantly to the costs they impose in the form of increased predation (Møller and Nielsen 1997; Zuk and Kolluru 1998; Candolin 2003; Godin and McDonough 2003; Johnson and Candolin 2017; but see Cain et al. 2019). Signals that are mediated by elevated testosterone can thus impose costs from both increased levels of the hormone and expression of the signal itself. Testosterone-mediated ornaments are often condition dependent (Bókony et al. 2008, McGlothlin et al. 2008), and thus these ornaments represent a potentially fruitful way to assess individual health in response to fluctuating environmental conditions (see Hill 1995). Most studies of the effect of perturbations on ornamental traits have been centered around degradation in response to pollution, most notably oil spills and mercury exposure, illustrating the potential for these signals to act as beacons for environmental health (Perez et al. 2010, Pérez et al. 2010, Jacques et al. 2019, Peneaux et al. 2020, Spickler et al. 2020). Determining whether natural disturbances like fire influence signaling traits, and the physiological mechanisms underlying these effects, holds promise both for informing conservation and understanding limits of phenotypic plasticity in disturbance-adapted organisms.

Fairywrens (family Maluridae) have emerged as model organisms for studies of the causes and consequences of male ornamentation. Young males of some Fairywren species undergo a pre-alternate molt prior to the breeding season in which most remain in drab female-like plumage whereas others molt into a showy ornamental plumage (Peters 2007, Lindsay et al. 2011). In Red-backed Fairywrens (*Malurus melanocephalus*), males adopting the sexually-selected ornamented phenotype circulate significantly higher testosterone than unornamented males (Lindsay et al. 2009) and testosterone-implantation induces molt into ornamental plumage (Lindsay et al. 2011, Khalil et al. 2020). Molt into ornamental plumage among young males seems to be condition-dependent, as experimentally-reduced condition caused second year males to remain unornamented for their first breeding season (Barron et al. 2013). Interestingly, young males in better condition acquired ornamental plumage absent of significantly greater testosterone circulation than males in worse condition (Barron et al. 2013), highlighting the need for additional studies to assess the intrinsic factors affecting ornament production in this system. In addition, resolving how extrinsic factors, including natural disturbances, influence expression of alternate phenotypes in this system could provide a useful test of condition-mediated signaling and its hormonal control in this system.

Here we capitalize on naturally occurring wildfires to determine effects on ornament expression of male Red-backed Fairywrens in tropical NE Queensland. We first determine whether a major wildfire prior to the typical breeding season in one of our long-term study populations affects molt into ornamental plumage. Next, we test whether known physiological mediators of ornamentation in this system, namely testosterone and fat stores (Lindsay et al. 2011, Barron et al. 2013), decrease shortly after wildfire. In addition, we determine whether corticosterone, a glucocorticoid hormone often associated with avian response to environmental stressors (Wingfield et al. 1998, Lucas et al. 2006, Bize et al. 2010, Busch et al. 2010, Wingfield 2013, Wingfield et al. 2017a) is elevated in response to fire. Finally, we assess what affects acquisition of ornamentation to reveal how prospective physiological mediators (fat, testosterone and corticosterone) shape alternate phenotype expression in response to a major environmental disturbance.

## METHODS

### Study system and timeline

The Red-backed Fairywren (*Malurus melanocephalus*) is a small, sedentary, insectivore of Australian savannas. Breeding occurs seasonally, with onset varying from year to year depending on monsoon rains (Webster et al. 2010). Males in their second year adopt one of two plumage phenotypes for breeding: brown (hereafter: unornamented) or black-and-red (hereafter: ornamented), and in rarer cases an intermediate mix of the two (Karubian et al. 2008, Webster et al. 2008). For the present study we employed plumage data from three separate years at our long-term color-banded “Donkey Farm” population (145°23’ E, 17°23’ S), and one year at a separate newly-banded “Sandy Creek” population (145°23’ E, 17°28’ S), both near Herberton, QLD, Australia. On October 12, 2012 a large wildfire burned our Donkey Farm field site when Fairywrens were beginning to form pairs prior to the typical breeding season in this population. The Donkey Farm fire burned the entire understory at our study site, leaving only a donkey paddock with short (<0.75m) grass that Fairywrens typically used sparingly during the non-breeding season. Following this fire, we established nearby (∼4km) Sandy Creek as a control site for determining acute effects of wildfire on fairy-wren physiology. On November 24 this site was also burned by a wildfire, providing natural replication of our experiment. The Sandy Creek fire also burned the vast majority of the understory at the site, but left several unburned patches smaller than a typical Fairywren breeding territory, and a horse paddock that was unoccupied by Fairywrens prior to the fire. Fairywrens in both populations were seen flocking at unusually high numbers in unburned patches, returning to territories at times to forage in the unburned *Eucalyptus* canopy and near unburned or regenerating *Lantana* shrubs. Individuals were caught at both populations in unburned grass and within recently burned habitat near *Lantana* shrubs and short *Eucalyptus* that survived fire. Both fires burned the sites in <24 hours, and we resumed sampling of Fairywrens 13 days after the Donkey Farm fire and 7 days after the Sandy Creek fire.

### Plumage assessment

Percent of ornamental black-and-red plumage (0-100) was assessed both at each capture of males and through continuous sightings of color-banded individuals from August to early January each year. We compared males exposed to fire in 2012 to previous seasons at our long-term Donkey Farm study site due to population-specific variation in ornamentation. Two previous field seasons were employed for this purpose: 2009-10, which was characterized by drought and minimal breeding but no fire, and 2012-13, which experienced a typical monsoon and breeding season for this population. This approach allowed us to assess both how fire affected acquisition of ornamental plumage relative to a similarly dry year with minimal breeding, and relative to a year with more typical rainfall and breeding.

### Field measurements

Red-backed Fairywrens were caught in mist nets by flushing or with use of playback of in-hand distress calls. We assessed size of furcular fat stores as a measure of condition by half increments from 0-3; 0 reflecting no fat and three representing fat protruding beyond the furcular hollow (Barron et al. 2013). Blood was taken immediately upon extraction of the bird from mist nets via the jugular vein (net to bleed time: 1.28 – 4.85 min.), then samples were placed in a small cooler for the remainder of the day. Upon return from the field, we spun blood samples in a centrifuge to separate plasma for hormone analysis. Plasma was stored in liquid nitrogen until shipment to the US where it was stored in a −20° freezer until use in hormone assays. Red blood cells were transferred to 400 µl of lysis buffer and refrigerated until use in genetic assays.

### Testosterone and corticosterone assay

We used a previously validated radioimmunoassay to measure both testosterone and corticosterone simultaneously for 13 to 50 µl plasma samples following Lindsay et al. (2011) and Barron et al. (2013). Briefly, we used tritium-labeled testosterone (Perkin Elmer Life Sciences NET-553) with a testosterone antibody (Wien Laboratories T-3003) that cross-reacts with other steroids (100% reactivity with testosterone, 60% with 5α-DHT, 5% aldosterone, <15% with other androgens, and less than 0.05% with 17β-estradiol and all other steroids), and tritium-labeled corticosterone with a corticosterone antibody (Esoterix Endocrinology B3-163). Samples were randomly assigned to 7 separate assays; the between-assay coefficient of variation was 12.96% for testosterone and 14.19% for corticosterone (calculated according to Chard 1995). The detection limit for testosterone was 276.32 pg/ml, and corticosterone was 0.81 ng/ml. We back-calculated undetectable testosterone levels from minimal detectable levels (1.95 pg/tube); all samples had detectable corticosterone. Testosterone in our dataset ranged from 76.56 to 8642.45 pg/ml (mean = 433.61 pg/ml, median = 424.19 pg/ml), and corticosterone was 0.97 to 24.61 ng/ml (mean = 6.36 ng/ml, median = 5.39 ng/ml).

### Sexing methods

Because unornamented males could not be sexed morphologically, we determined sex genetically. We extracted DNA from blood samples using the Omega Bio-Tek EZ 96 Total DNA/RNA Isolation Kit® and amplified an intron within the CHD gene using the primer pair 1237L/1272H (Kahn et al. 1998). Briefly, we PCR-amplified each sample following the protocol of Varian-Ramos et al. (2010), and visualized the PCR products via electrophoresis on a 2% agarose gel. We ran samples alongside a positive and negative control, and visually inspected the gels. We considered individuals with a single band to be males, and individuals with two bands to be females.

### Data analysis

All analyses were conducted in R studio version 1.2.5019 (R Core Team 2018) using nonparametric tests in base R, and linear models in base R and package lme4 (Bates et al. 2015).

#### Effect of wildfire on phenotype expression

We filtered data from each year at our Donkey Farm study site (2009, 2011, and 2012) to analyze males who were of known age. We only analyzed males who were observed after December 1^st^ each year, as males have typically assumed their final plumage phenotype well before this date in our study population (Barron, pers. comm.). Males known to be in their second and third year, as well as males known to be older than third year were analyzed individually in separate models to determine which age classes contributed to population-wide differences in ornament expression. We used a binomial mixed model for analysis of all known age males. Age and date of final plumage observation were included as fixed effects, and individual ID was included as a random effect in this model to account for repeated measures. We used nonparametric Kruskal Wallis tests for assessing differences in final plumage by year among discrete age classes. Following detection of significant differences in phenotype expression we used post hoc Tukey comparisons for the mixed model containing all males, and pair-wise Wicoxon Rank Sum test with Holm-Bonferroni correction for discrete age classes to determine which years differed.

#### Effect of wildfire on physiological mediators of phenotype expression

We pooled capture data from the Donkey Farm and Sandy Creek populations, who were exposed to separate wildfires 33 days apart, which resulted in natural replication of our experiment. Due to lack of demographic information from the Sandy Creek population we were unable to account for potential age effects. However, by filtering both populations to only include males who were initially observed with <33% ornamental plumage we were able to remove age classes that were unable to mount a plastic response to environmental conditions. We used this data set to determine how furcular fat stores, and plasma testosterone and corticosterone levels were affected by fire among males who were were sampled within 30 days before and after onset of fire. Furcular fat stores and testosterone were log transformed, and corticosterone levels were square root transformed to improve normality. For hormone models we first calculated residuals based on models that included time of day bled, capture to bleed time, and day of year as fixed effects according to Vernasco et al. (2019). We then analyzed all physiological components using linear models. Population and the interaction between population and treatment (pre vs. post fire) were fixed effects in all models, and individual ID a random effect to control for repeated measures. Finally, we assessed whether circulating testosterone and corticosterone were correlated with a linear regression.

#### What explains variation in phenotype expression?

We compiled post-fire data from both populations exposed to wildfire in 2012 and selected males who were first observed as brown and were actively molting at the time of recapture. We analyzed the effect of log transformed furcular fat stores and testosterone and corticosterone residuals calculated according to above methods on final plumage ornamentation score. Males who were not observed after December 1^st^ were excluded from analyses, and date of final observation and site were included as fixed effects. We used model selection to choose the best regression model by AIC due to an absence of an *a priori* hypothesis about the mechanism(s) underlying diminished ornament expression. Additionally, we analyzed whether males molting in exclusively unornamented or ornamented plumage differed according to significant predictor(s) of final phenotype expression revealed in our initial regression models.

## RESULTS

### Effect of wildfire on phenotype expression

There was a significant effect of year (dry, wet, and fire year) and age on final plumage ornamentation score (Figure 2) when all age classes were pooled (Year: *W*^*2*^(2, *N* = 148) = 14.19, *p* = 0.00083; Age: *W*^*2*^(1, *N* = 148) = 31.32, *p* < 0.0001). A post-hoc Tukey test revealed that males were significantly less ornamented in the fire year compared to the dry year (i.e. drought conditions; *p* = 0.0018) and wet year (i.e. typical monsoon conditions; *p* = 0.0038) while final ornamentation did not differ between the wet and dry year (*p* = 0.15). Analysis of known second-year males showed significantly less ornamentation in the fire year relative to the wet year (H(1) = 14.28, *p* = 0.00016); limited sample sizes precluded inclusion of the dry year in this analysis. Among third-year males ornamentation also differed across years (H(2) = 15.5, *p* = 0.00043); a pairwise post-hoc Wilcoxon test revealed that third-year males during the fire year were significantly less ornamented relative both to the dry (*p* = 0.0064) and wet year (*p* = 0.0037), while ornamentation did not differ between the dry and wet year (*p* = 0.15). Final ornamentation score did not differ across years in pooled older age classes (>3^rd^ year; H(2) = 3.006, *p* = 0.22). Date of final plumage observation was not a significant predictor in any models.

**Figure 1.**
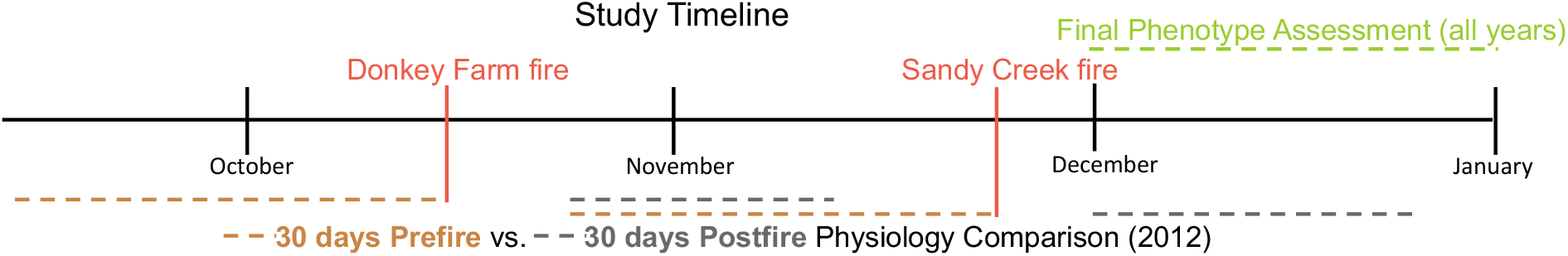
Study timeline, depicting when fires occurred at both Donkey Farm and Sandy Creek populations (near Herberton, QLD, Australia. Physiological sampling occurred 30 days prior to and 13-30 days after the 2012 Donkey Farm fire. We sampled Sandy Creek 30 days prior to and 7-30 days after it burned in a separate fire. Final phenotype assessment was conducted after December 1^st^ in all study years (2009, 2011, 2012).

**Figure 2.**
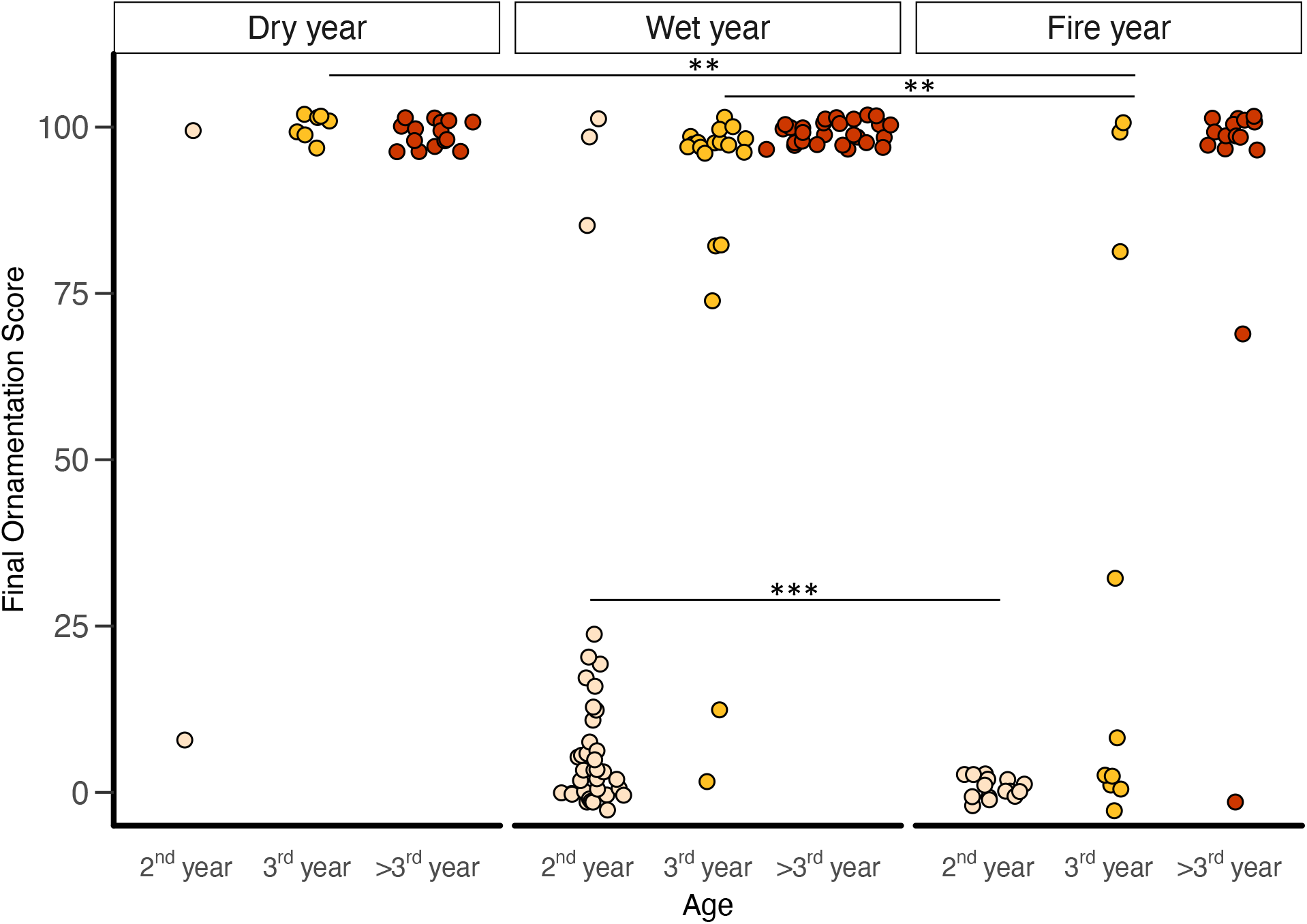
Comparison of final plumage ornamentation (score 0 = fully brown plumage; score 100 = fully red/black plumage) across three study years varying in environmental conditions (dry, wet, fire year) at the Donkey Farm study population. Males in the fire year were less ornamented when all age classes were pooled; 2^nd^ and 3^rd^ year males were less ornamented when comparing the fire year to the wet and dry year. Limited sample sizes precluded comparison of 2^nd^ year males from the dry year to other years. Significant differences between years are indicated with asterisks (\*\**p* < 0.01, \*\*\**p* < 0.001).

### Effect of wildfire on physiological mediators of phenotype expression

We compared physiological measures collected within 30 days prior to and 30 days post fire from both populations exposed to wildfire (Table 1). For the comparison of hormone levels we used residuals from regression against time of day and time to obtain blood samples which affect hormone levels (see Methods). We did not find significant differences in furcular fat stores (F_1, 60.78_ = 0.11, *p* = 0.74) or testosterone residuals (F_1, 20.06_ = 0.67, *p* = 0.42). Corticosterone residuals were significantly higher after fire (F_1, 35.01_ = 6.02, *p* = 0.019; Figure 3). None of the potential mediators differed by population (fat: F_1, 62.62_ = 0.14, *p* = 0.71, testosterone residuals: F_1, 47.52_ = 1.77, *p* = 0.19, corticosterone residuals: F_1, 37.05_ = 0.0021, *p* = 0.96). The interaction between condition (pre-versus post-fire) and population was significant for fat (F_1, 60.78_ = 5.31, *p* = 0.025), but not testosterone (F_1, 20.06_ = 0.06, *p* = 0.81) or corticosterone (F_1, 35.01_ = 0.73, *p* = 0.40) residuals.

**Table 1.**
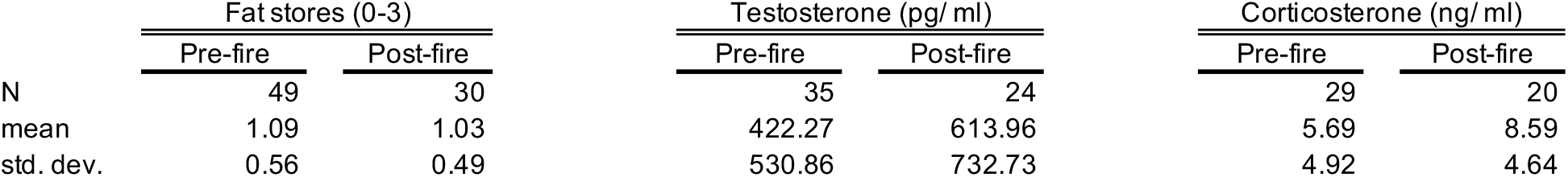
Summary statistics for fat stores, circulating corticosterone, and circulating testosterone for individuals sampled within 30 days pre-fire and 30 days post-fire across two populations.

**Figure 3.**
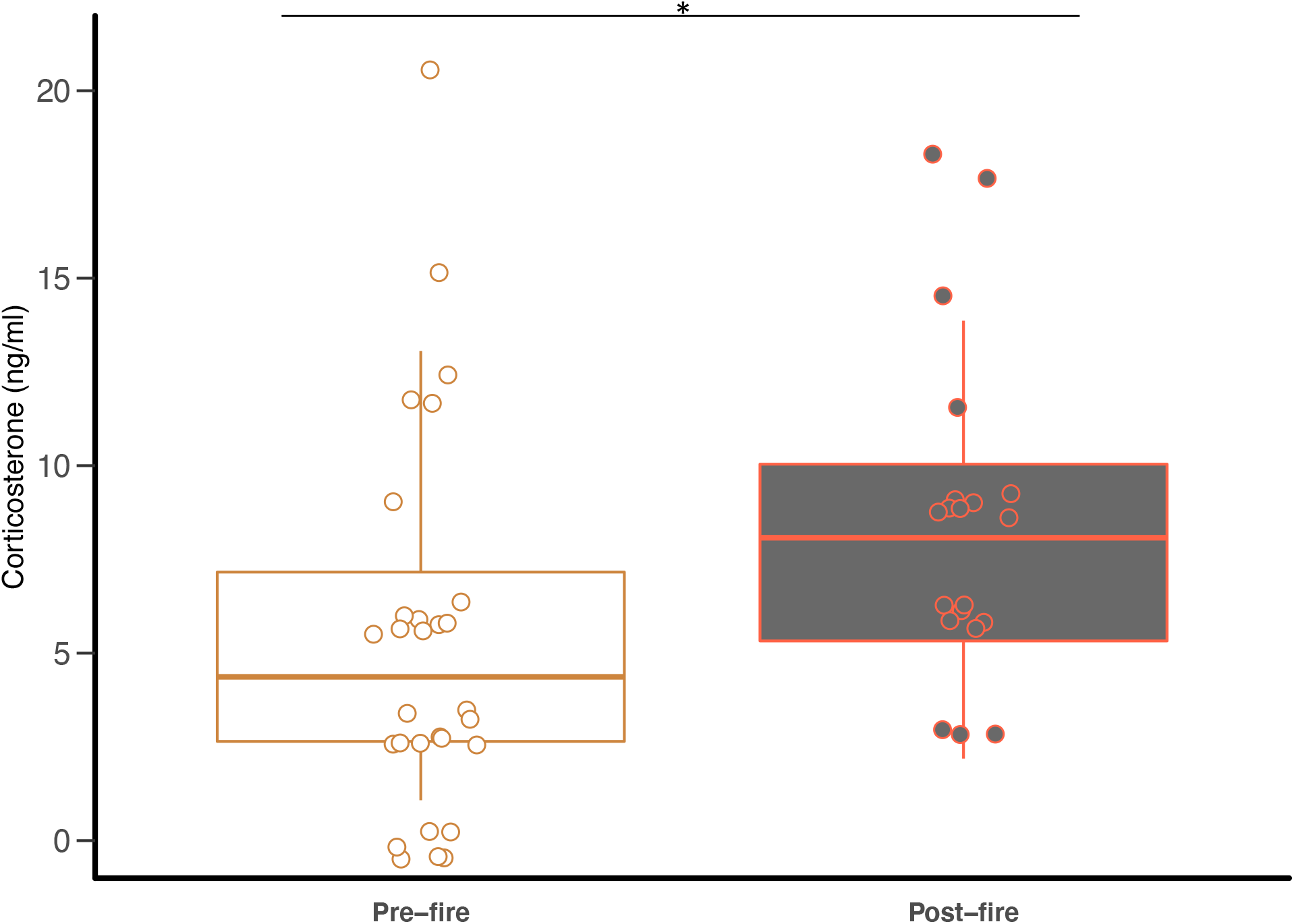
Circulating corticosterone (ng/ ml) for males sampled within 30 days prior to (“Pre-fire”) and 30 days after (“Post-fire”) wildfires, pooling across two sites. Box-and-whisker plot depicts the median, first quartile and third quartile, with individual data points overlaid. Corticosterone residuals (accounting for capture time and capture to bleed time) were significantly higher after fire (*p* < 0.05).

### What explains variation in phenotype expression?

The best model explaining final ornamentation scores across both sites exposed to wildfire included testosterone residuals (F_1, 43.80_ = 15.66, *p* = 0.0032) and population (F_1, 18.74_ = 6.70, *p* = 0.014) as significant predictors and date of final observation as a non-signficant factor (F_1, 1.54_ = 1.0, *p* = 0.33). Testosterone was significantly higher among males molting in exclusively ornamental red and black feathers vs. males molting exclusively brown feathers (χ^2^ (1, N = 37) = 8.08, *p* = 0.0045; Figure 4), but not by population (χ^2^ (1, N = 37) = 0.75, *p* = 0.39). There was no correlation between circulating testosterone and corticosterone residuals (R^2^ = 0.0085, F_1,119_=1.01, *p* = 0.32; Figure S1).

**Figure 4.**
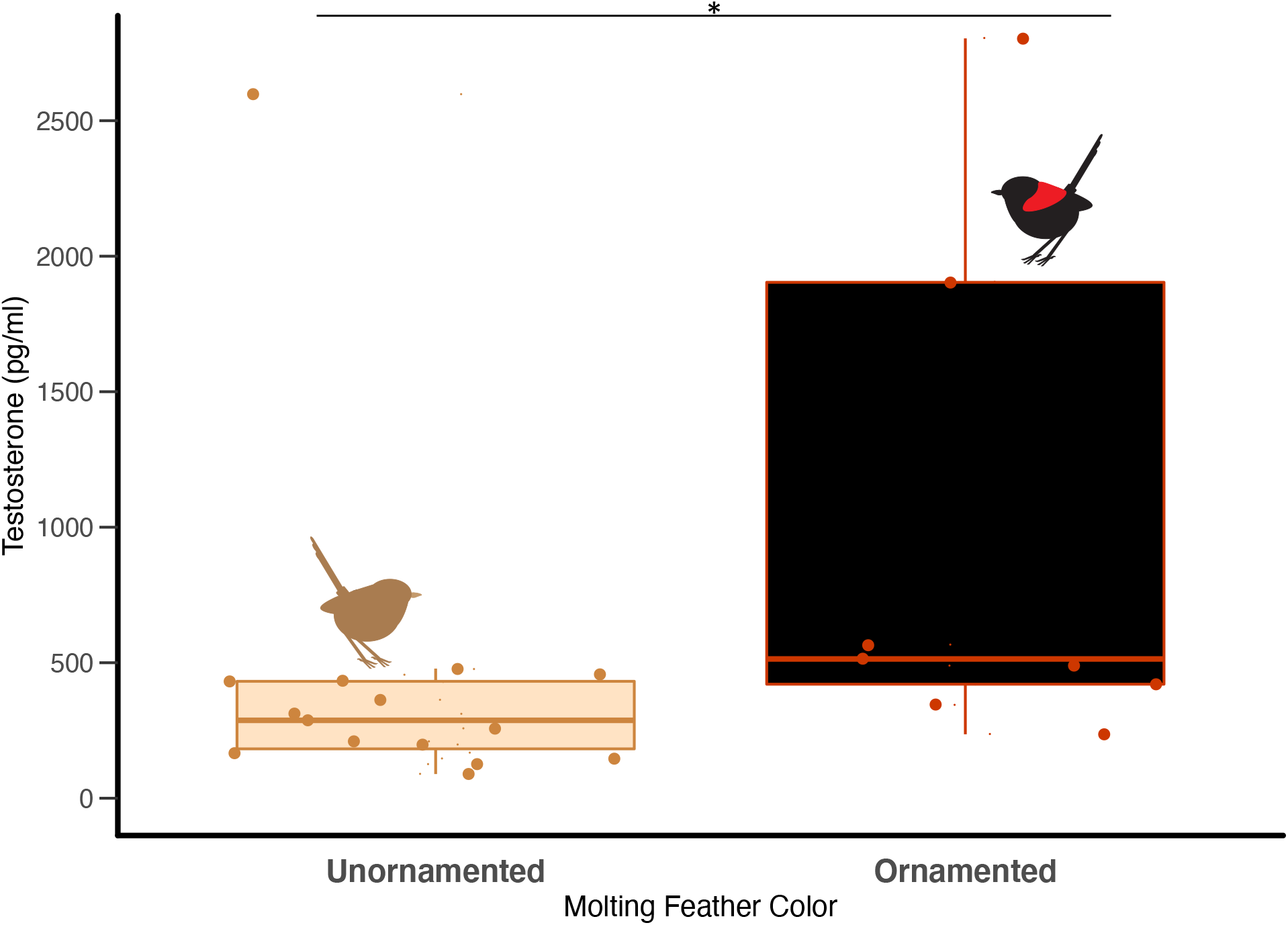
Circulating testosterone (pg/ml) in males molting in exclusively unornamented (brown/white) vs. ornamented (red/black) plumage (*p* = 0.019). As in Figure 3, box-and-whisker depicts the median, first quartile and third quartile, with individual data points overlaid (excluding a single outlier).

## DISCUSSION

We found that a wildfire affected phenotype expression in male Red-backed Fairywrens by decreasing the proportion of young males molting into the sexually-selected, ornamented red/black plumage (Figure 2). Surprisingly, males in their third year also exhibited substantial variation in extent of ornamental plumage, despite previous studies indicating relatively little plasticity in this age class (Webster et al. 2008). Although we did not find that testosterone decreased shortly after wildfires in males starting the year in brown plumage (Table 1), the importance of testosterone for ornament production in this species is supported by a positive correlation between testosterone and ornamentation, and by the fact that males molting into exclusively ornamented plumage exhibited higher testosterone levels than did males molting into drab plumage (Figure 4). Furcular fat stores and baseline plasma testosterone levels did not change shortly (<30 days) after fire, indicating that these physiological mediators aren’t especially sensitive to this type of disturbance in the short-term. In contrast, elevated baseline corticosterone (Figure 3) in males recently exposed to fire could be indicative of short-term stress. However, neither fat stores nor corticosterone levels were predictive of final ornamentation score, so diminished condition or higher corticosterone was unlikely to have caused suppressed plumage ornamentation following fire.

The current study clarifies the extent of delayed plumage maturation in Red-backed Fairywrens and the role of previously identified physiological mediators underlying plasticity in phenotype expression. Previous studies on delayed plumage maturation in this system have addressed causes and consequences, focusing on second year males in their first breeding season due to perceived lack of plasticity in older males (Karubian et al. 2008, Webster et al. 2008). Correlational studies (Lindsay et al. 2009) as well as manipulations of testosterone (Lindsay et al. 2011) and body condition (Barron et al. 2013) in second year males established both as mediators of alternate plumage phenotype expression. Specifically, testosterone levels were higher in males molting into red/black ornamented plumage compared to males that molted into cryptic brown, female-like plumage during both the pre-breeding and the reproductive period (Lindsay et al. 2009). In separate experiments, testosterone implants induced molt of ornamented plumage (Lindsay et al. 2011; Khalil et al. 2020); and, increased body condition in response to experimental manipulation resulted in more ornamentation, however, without an associated increase in circulating testosterone (Barron et al. 2013). Adding further complexity to the relationship between testosterone and ornamentation in this system, older, early molting males seem to acquire ornamented plumage absent of an elevation in testosterone (Lantz et al. 2017).

Results from the present study support a complex role for testosterone in plumage ornamentation. In addition, we reveal greater plasticity in phenotype expression in this system than previously assumed: males in their third year exhibited uncharacteristically diminished ornament expression during the wildfire year (Figure 2). Importantly, these males were significantly less ornamented during the fire year when compared to the same age category in a typical year with seasonal monsoon rain, and in a drought year without fire. There are multiple interpretations of causality for this key result. First, third year males exposed to drought ready themselves for breeding (i.e. produce sexually-selected plumage) in anticipation of seasonally variable monsoon rain (Webster et al. 2010), whereas fire destroys nesting substrate for several weeks to months and thus precludes the need to prepare for breeding. Second, third year males could have been limited in their ability to enhance testosterone circulation and body condition required for ornament production due to limited foraging substrate following fire. Limited physiological sampling from known third year males precluded our ability to assess whether testosterone or body condition metrics were diminished following fire in these males. However, when pooling males who started the year unornamented across two sites exposed to fire, we did not find a short-term (within 30 days) decrease in plasma testosterone levels or reduction of fat stores (Table 1). Given that both fires occurred when males typically elevate testosterone to molt into ornamental plumage (Lindsay et al. 2009), our null result in pre-fire vs. post-fire comparisons could be indicative of a shift from typical seasonal patterns of testosterone circulation. We found that testosterone levels following fire were predictive of final plumage phenotype, and that males molting exclusively ornamental plumage had higher plasma testosterone than those molting exclusively unornamented plumage (Figure 4). Overall, our results indicate that diminished ornamentation following fire is at least partially the result of subtle effects to testosterone circulation.

Glucocorticoids have been proposed as mediators of the tradeoff between survival and reproduction in vertebrates (Breuner et al. 2008, Bonier et al. 2009, Almasi et al. 2013, Crespi et al. 2013). Elevated corticosterone can initiate an emergency life history stage to flee from a natural disturbance, thus interrupting typical phenology (Wingfield et al. 1998, Wingfield and Kitaysky 2002). Many studies of bird species have revealed that corticosterone is responsive to climatic events, and can mediate migration away from breeding territories when conditions deteriorate (Astheimer et al. 1995, Wingfield and Kitaysky 2002, Lynn et al. 2003, Wingfield 2013, Krause et al. 2016, Wingfield et al. 2017b). Additionally, one way in which baseline glucocorticoids are thought to mediate the tradeoff between survival and mating is through antagonistic effects on testosterone (Stanczyk et al. 1985, McGuire et al. 2013). In our study, corticosterone was elevated in males sampled 7-30 days after fire relative to males sampled within 30 days before fire. Our post-fire sampling window corresponded to a time when Red-backed Fairywrens were off pair-defended territories foraging in large flocks in unburned paddocks and small patches of unburnt grassland at both populations, consistent with a role of elevated corticosterone in triggereing changes in behavior. However, absolute differences in baseline plasma corticosterone are small enough (Table 1; Figure 3) to warrant caution in interpreting our results as indicative of a corticosterone-induced emergency life history stage. We did not resume sampling until 13 days after the Donkey Farm fire and 7 days after the Sandy Creek fire, which precluded our ability to measure a truly acute (i.e. within 24 hours) response to these events. In addition, this limited sampling window could have kept us from detecting diminished fat stores following fire (Table 1). Finally, we did not find a relationship between corticosterone and testosterone in our dataset (Figure S1), thus adding to a growing list of studies in birds suggesting an absence of antagonism between these two hormones (Davies et al. 2016, Deviche et al. 2017, Abolins-Abols et al. 2018).

Species that evolved in unpredictable environments characterized by regular natural disturbance (e.g. drought, fires) might reasonably be expected to mount an adaptive response to perturbation. Australian tropical savannas are characterized by a history of frequent wildfires, but many species inhabiting these areas are in decline and are negatively-affected by fire (Franklin et al. 2005, Woinarski and Legge 2013). Previous studies of Red-backed Fairywrens in this context have shown that wildfires lead to decreased reproductive output (Murphy et al. 2010), increased social connectivity (Lantz and Karubian 2017) and shifting habitat use (Sommer et al. 2018) in this species. Although our study did not focus on these characteristics, our observations of Red-backed fairywrens support these findings as wildfires forced individuals to leave breeding territories and flock in unusual numbers in the few remaining unburned areas at both study sites (Boersma, pers. obs.). A companion study to ours found reduced dawn singing following wildfire, indicative of a reduction in behavior associated with breeding (Mathers-Winn et al. 2018). We found just one active nest during the study period, which extended through the typical breeding season at our long-term study site, so our findings are consistent with a major reduction or delay in breeding effort following wildfires (Boersma, pers. obs.) as previously reported in this species (Murphy et al. 2010). Our results suggest that Red-backed Fairywrens exposed to fire prior to the breeding season delay acquisition of potentially costly sexual signals through molt, and thus perhaps buffer against detrimental costs to survival.

## Conclusion

Wildfires affected young male Red-backed Fairywrens by suppressing or greatly delaying molt into ornamental breeding plumage. While corticosterone was elevated in the short-term, an important metric of body condition in these birds (furcular fat stores) was unaffected by fire. Circulating testosterone after fire was predictive of final plumage ornamentation score in young males, consistent with the previously established role for testosterone in production of ornamentation in this system. These results are consistent with this species acting to maximize survival through changes in hormones when a natural disturbance destroys breeding territories. As the prevalence of wildfires increases, it is important to understand how and to which extent organisms can acutely cope to improve predictive models of response to climate change across diverse taxa. Most studies of response to wildfire have focused on broad and long-term effects to survival and reproduction. More research is needed to determine how physiological mechanisms underlying the tradeoff between survival and reproduction are affected by fire and other natural disturbances, as presented here.

## DECLARATIONS

## Acknowledgements

We are grateful to T. Belluz, E. Fishel, J. Hiciano, C. Mathers-Winn, and E. Williams for expert assistance in the field in 2012-13, and to field crews from 2009-10 and 2011-12. J. Dowling provided indispensable guidance and T. Daniel provided vital logistical support in the field. P. Carter, E. Crespi, and H. Watts and B. Vernasco provided valuable advice on data analysis and manuscript revision. S. Khalil contributed Fairywren icons used in plumage figures.

## Funding

This study was supported by a National Science Foundation Grant (#0818962) to MSW and HS.

## Author contribution

JB and DGB devised the project; JB collected data; HS and MSW provided funding and helped with experimental design; JB did endocrine lab work and DTB did genetic work; JB analyzed data; JB wrote manuscript; all authors helped revise the paper.

## Conflicts of interest

The authors declare no conflict of interest.

## Permits

Permission for this work was granted by the Queensland Government Encironmental Protection Agency. Field protocols were approved by the Institutional Animal Care and Use Committee of Washington State University. Samples were exported from Australia with approval by the Australian Government Department of Environment and Heritage.

## SUPPLEMENT

**Figure S1.**
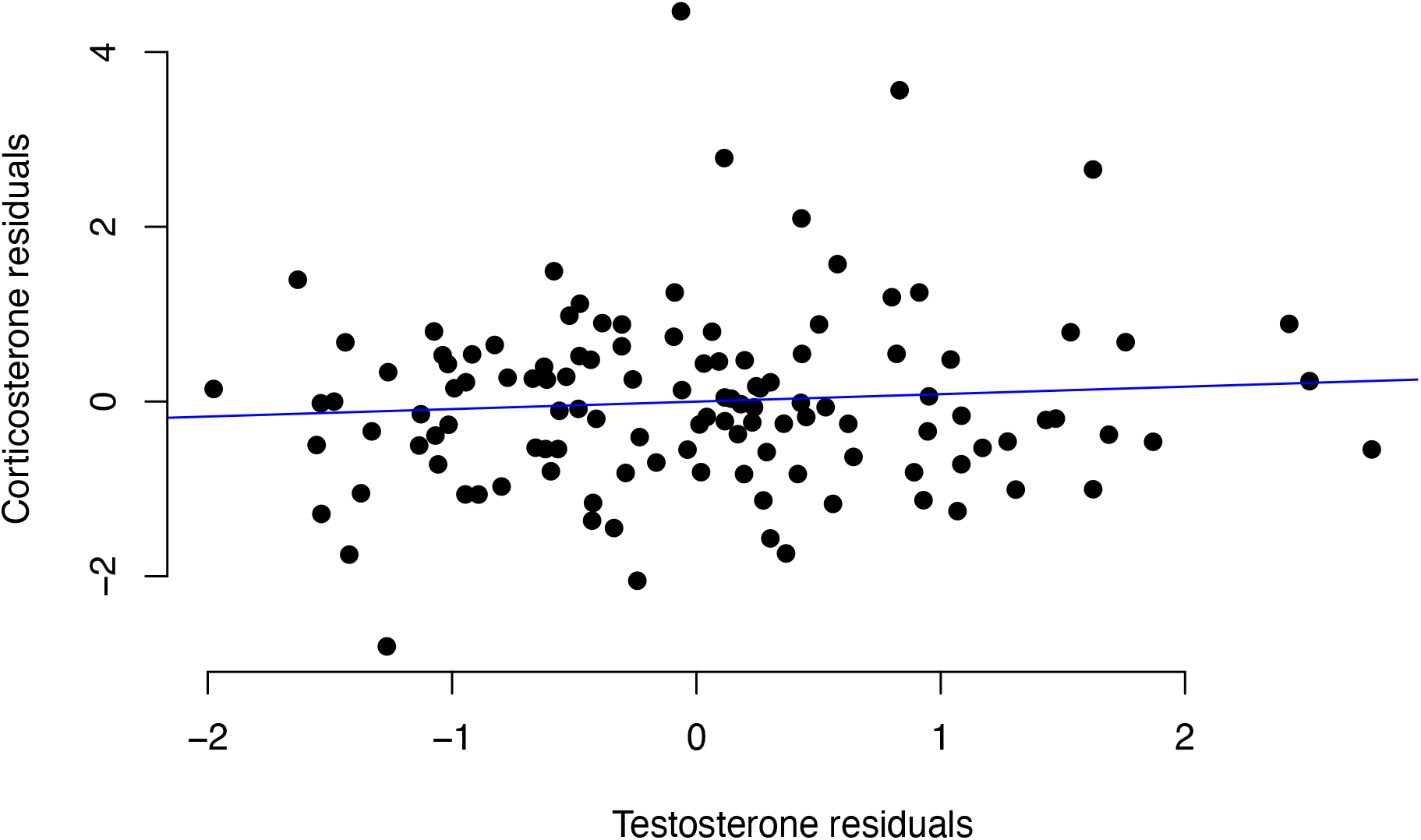
Correlation between log transformed baseline testosterone and square root transformed baseline corticosterone following exposure to fire.

## Notes

### Competing Interest Statement

The authors have declared no competing interest.

## REFERENCES

Abolins-Abols, M., Hanauer, R. E., Rosvall, K. A., Peterson, M. P. and Ketterson, E. D. 2018. The effect of chronic and acute stressors, and their interaction, on testes function: An experimental test during testicular recrudescence. - J. Exp. Biol. 221(17): jeb180869.

Almasi, B., Roulin, A. and Jenni, L. 2013. Corticosterone shifts reproductive behaviour towards self-maintenance in the barn owl and is linked to melanin-based coloration in females. - Horm. Behav. 64: 161–171.

Astheimer, L. B., Buttemer, W. A. and Wingfield, J. C. 1995. Seasonal and acute changes in adrenocortical responsiveness in an arctic-breeding bird. - Horm. Behav. 29: 442–457.

Barlow, J. and Peres, C. A. 2004a. Avifaunal responses to single and recurrent wildfires in Amazonian forests. -Ecol. Appl. 14: 1358–1373.

Barlow, J. and Peres, C. A. 2004b. Ecological responses to El Niño-induced surface fires in central Brazilian Amazonia: Management implications for flammable tropical forests. - Philos. Trans. R. Soc. B Biol. Sci. 359: 367–380.

Barlow, J., Haugaasen, T. and Peres, C. A. 2002. Effects of ground fires on understorey bird assemblages in Amazonian forests. - Biol. Conserv. 105: 157–169.

Barlow, J., Peres, C. A., Henriques, L. M. P., Stouffer, P. C. and Wunderle, J. M. 2006. The responses of understorey birds to forest fragmentation, logging and wildfires: An Amazonian synthesis. - Biol. Conserv. 128: 182–192.

Barron, D. G., Webster, M. S. and Schwabl, H. 2013. Body condition influences sexual signal expression independent of circulating androgens in male red-backed fairy-wrens. - Gen. Comp. Endocrinol. 183: 38–43.

Bates, D., Maechler, M., Bolker, B. and Walker, S. 2015. Fitting Linear Mixed-Effects Models Using lme4. - J. Stat. Softw. 67: 1–48.

Bize, P., Stocker, A., Jenni-Eiermann, S., Gasparini, J. and Roulin, A. 2010. Sudden weather deterioration but not brood size affects baseline corticosterone levels in nestling Alpine swifts. - Horm. Behav. 58: 591–598.

Bókony, V., Garamszegi, L. Z., Hirschenhauser, K. and Liker, A. 2008. Testosterone and melanin-based black plumage coloration: A comparative study. - Behav. Ecol. Sociobiol. 62: 1229–1238.

Bonier, F., Martin, P. R., Moore, I. T. and Wingfield, J. C. 2009. Do baseline glucocorticoids predict fitness? - Trends Ecol. Evol. 24: 634–642.

Breuner, C. W., Patterson, S. H. and Hahn, T. P. 2008. In search of relationships between the acute adrenocortical response and fitness. - Gen. Comp. Endocrinol. 157: 288–295.

Busch, D. S., Addis, E. A., Clark, A. D. and Wingfield, J. C. 2010. Disentangling the effects of environment and life-history stage on corticosterone modulation in costa rican rufous-collared sparrows, Zonotrichia capensis costaricensis. - Physiol. Biochem. Zool. 83: 87–96.

Cain, K. E., Hall, M. L., Medina, I., Leitao, A. V, Delhey, K., Brouwer, L., Peters, A., Pruett-jones, S., Webster, M. S., Langmore, N. E. and Mulder, R. A. 2019. Conspicuous Plumage Does Not Increase Predation Risk?: A Continent-Wide Test Using Model Songbirds. - Am. Nat. 193(3): 359–372.

Candolin, U. 2003. The use of multiple cues in mate choice. - Biol. Rev. 78: 575–595.

Cawthorn, J. M., Morris, D. L., Ketterson, E. D. and Nolan, V. 1998. Influence of experimentally elevated testosterone on nest defence in dark-eyed juncos. - Anim. Behav. 56: 617–621.

Chard, T. 1995. An Introduction to Radioimmunoassay and Related Techniques. Laboratory Techniques in Biochemistry and Molecular Biology. - Elsevier: Oxford.

Cook, B. I., Smerdon, J. E., Seager, R. and Coats, S. 2014. Global warming and 21st century drying. - Clim. Dyn. 43: 2607–2627.

Cox, J. A. and Widener, B. 2008. Lightning-Season Burning: Friend or Foe of Breeding Birds? - Tall Timbers Res. Stn. Misc. Publ. 17: 1–16.

Crespi, E. J., Williams, T. D., Jessop, T. S. and Delehanty, B. 2013. Life history and the ecology of stress: How do glucocorticoid hormones influence life-history variation in animals? - Funct. Ecol. 27: 93–106.

Dai, A. 2013. Increasing drought under global warming in observations and models. - Nat. Clim. Chang. 3: 52–58.

Davies, S., Noor, S., Carpentier, E. and Deviche, P. 2016. Innate immunity and testosterone rapidly respond to acute stress, but is corticosterone at the helm? - J. Comp. Physiol. B Biochem. Syst. Environ. Physiol. 186: 907–918.

Deviche, P., Desaivre, S. and Giraudeah, M. 2017. Experimental manipulation of corticosterone does not influence the clearance rate of plasma testosterone in birds. - Physiol. Biochem. Zool. 90: 575–582.

Flannigan, M., Cantin, A. S., De Groot, W. J., Wotton, M., Newbery, A. and Gowman, L. M. 2013. Global wildland fire season severity in the 21st century. - For. Ecol. Manage. 294: 54–61.

Foo, Y. Z., Nakagawa, S., Rhodes, G. and Simmons, L. W. 2017. The effects of sex hormones on immune function: A meta-analysis. - Biol. Rev. 92: 551–571.

Franklin, D. C., Whitehead, P. J., Pardon, G., Matthews, J., McMahon, P. and McIntyre, D. 2005. Geographic patterns and correlates of the decline of granivorous birds in northern Australia. - Wildl. Res. 32: 399–408.

Godin, J. G. J. and McDonough, H. E. 2003. Predator preference for brightly colored males in the guppy: A viability cost for a sexually selected trait. - Behav. Ecol. 14: 194–200.

Hau, M. 2007. Regulation of male traits by testosterone: Implications for the evolution of vertebrate life histories. - BioEssays 29: 133–144.

Hill, G. E. 1995. Ornamental traits as indicators of environmental health. - Bioscience 45: 25–30.

Hobson, K. A. and Schieck, J. 1999. Changes in bird communities in boreal mixedwood forest: Harvest and wildfire effects over 30 years. - Ecol. Appl. 9: 849–863.

Jacques, D. L., Mills, K. L., Selby, B. G., Veit, R. R. and Id, H. Z. 2019. Use of plumage and gular pouch color to evaluate condition of oil spill rehabilitated California brown pelicans (Pelecanus occidentalis californicus) post-release. - PLoS One 142: e0211932.

Johnson, S. and Candolin, U. 2017. Predation cost of a sexual signal in the threespine stickleback. - Behav. Ecol. 28: 1160–1165.

Kahn, N. W., St John, J. and Quinn, T. W. 1998. Chomosome-specific intron size differences in the avian CHD gene provide an efficient method for sex identification in birds. - Auk 115: 1074–1078.

Karubian, J., Sillett, T. S. and Webster, M. S. 2008. The effects of delayed plumage maturation on aggression and survival in male red-backed fairy-wrens. - Behav. Ecol. 19: 508–516.

Khalil, S., Welklin, J. F., McGraw, K. J., Boersma, J., Schwabl, H., Webster, M. S. and Karubian, J. 2020. Testosterone regulates CYP2J19-linked carotenoid signal expression in male red-backed fairywrens (Malurus melanocephalus): Testosterone and CYP2J19 expression. - Proc. R. Soc. B Biol. Sci. 287: 20201687.

Kimball, R. T. and Ligon, J. D. 1999. Evolution of avian plumage dichromatism from a proximate perspective. - Am. Nat. 154: 182–193.

Krause, J. S., Pérez, J. H., Chmura, H. E., Sweet, S. K., Meddle, S. L., Hunt, K. E., Gough, L., Boelman, N. and Wingfield, J. C. 2016. The effect of extreme spring weather on body condition and stress physiology in Lapland longspurs and white-crowned sparrows breeding in the Arctic. - Gen. Comp. Endocrinol. 237: 10–18.

Lantz, S. M. and Karubian, J. 2017. Environmental disturbance increases social connectivity in a passerine bird. - PLoS One 12: 1–15.

Lantz, S. M., Boersma, J., Schwabl, H. and Karubian, J. 2017. Early-moulting Red-backed Fairywren males acquire ornamented plumage in the absence of elevated androgens. - Emu - Austral Ornithol. 117(2): 170–180.

Lemos Da Silva, T., Lemes Marques, E. and Guilherme, E. 2015. Recuperation of the terra firme forest understory bird fauna eight years after a wildfire in eastern Acre, Brazil. - Int. J. Ecol.

Lindsay, W. R., Webster, M. S., Varian, C. W. and Schwabl, H. 2009. Plumage colour acquisition and behaviour are associated with androgens in a phenotypically plastic tropical bird. - Anim. Behav. 77: 1525–1532.

Lindsay, W. R., Webster, M. S. and Schwabl, H. 2011. Sexually selected male plumage color is testosterone dependent in a tropical passerine bird, the red-backed fairy-wren (Malurus melanocephalus). - PLoS One 6: e26067.

Lucas, J. R., Freeberg, T. M., Egbert, J. and Schwabl, H. 2006. Fecal corticosterone, body mass, and caching rates of Carolina chickadees (Poecile carolinensis) from disturbed and undisturbed sites. - Horm. Behav. 49: 634–643.

Lynn, S. E., Hunt, K. E. and Wingfield, J. C. 2003. Ecological factors affecting the adrenocortical response to stress in chestnut-collared and McCown’s longspurs (Calcarius ornatus, Calcarius mccownii). - Physiol. Biochem. Zool. 76: 566–576.

Martin, T. E. and Mouton, J. C. 2020. Longer-lived tropical songbirds reduce breeding activity as they buffer impacts of drought. - Nat. Clim. Chang. 10: 953–958.

Mathers-Winn, C., Dowling, J. L. and Webster, M. S. 2018. Forest fire reduces dawn singing effort in a passerine bird. - Aust. F. Ornithol. 35: 75.

McGlothlin, J. W., Jawor, J. M., Greives, T. J., Casto, J. M., Phillips, J. L. and Ketterson, E. D. 2008. Hormones and honest signals: Males with larger ornaments elevate testosterone more when challenged. - J. Evol. Biol. 21: 39–48.

McGlothlin, J. W., Whittaker, D. J., Schrock, S. E., Gerlach, N. M., Jawor, J. M., Snajdr, E. A. and Ketterson, E. D. 2010. Natural Selection on Testosterone Production in a Wild Songbird Population. - Am. Nat. 175: 687–701.

McGuire, N. L., Koh, A. and Bentley, G. E. 2013. The direct response of the gonads to cues of stress in a temperate songbird species is season-dependent. - PeerJ 1: e139.

Mestre, L. A. M., Cochrane, M. A. and Barlow, J. 2013. Long-term Changes in Bird Communities after Wildfires in the Central Brazilian Amazon. - Biotropica 45: 480– 488.

Møller, A. P. and Nielsen, J. T. 1997. Differential predation cost of a secondary sexual character: Sparrowhawk predation on barn swallows. - Anim. Behav. 54: 1545– 1551.

Murphy, S. A., Legge, S. M., Heathcote, J. and Mulder, E. 2010. The effects of early and late-season fires on mortality, dispersal, physiology and breeding of red-backed fairy-wrens (Malurus melanocephalus). - Wildl. Res. 37: 145–155.

Peneaux, C., Hansbro, P. M. and Griffin, A. S. 2020. The potential utility of carotenoid-based coloration as a biomonitor of environmental change. - Ibis: in press.

Perez, C., Munilla, I., López-Alonso, M. and Velando, A. 2010. Sublethal effects on seabirds after the Prestige oil-spill are mirrored in sexual signals. - Biol. Lett. 6: 33– 35.

Pérez, C., Lores, M., Velando, A., Pérez, C., Lores, M. and Velando, A. 2010. Oil pollution increases plasma antioxidants but reduces coloration in a seabird. - Oecologia 163: 875–884.

Peters, A. 2000. Testosterone treatment is immunosuppressive in superb fairy-wrens, yet free-living males with high testosterone are more immunocompetent. - Proc. Biol. Sci. 267: 883–889.

Peters, A. 2007. Testosterone treatment of female Superb Fairy-wrens Malurus cyaneus induces a male-like prenuptial moult, but no coloured plumage. - Ibis (Lond. 1859). 149: 121–127.

R Core Team 2018. R: A language and environment for statistical computing. in press.

Reed, W.L., Clark, M. E., Parker, P. G., Raouf, S. A., Arguedas, N., Monk, D. S.,Snajdr, E., Nolan, V. and Ketterson, E. D. 2006. Physiological effects on demography: A long-term experimental study of testosterone’s effects on fitness. - Am. Nat. 167: 667–683.

Roberts, M. L., Buchanan, K. L. and Evans, M. R. 2004. Testing the immunocompetence handicap hypothesis: A review of the evidence. - Anim. Behav. 68: 227–239.

Şekercioĝlu, çaĝan H., Primack, R. B. and Wormworth, J. 2012. The effects of climate change on tropical birds. - Biol. Conserv. 148: 1–18.

Sommer, N. R., Moody, N. M., Lantz, S. M., Leu, M., Karubian, J. and Swaddle, J. P. 2018. Red-backed fairywrens adjust habitat use in response to dry season fires. - Austral Ecol. 43: 876–889.

Spickler, J. L., Swaddle, J. P., Gilson, R. L. and Daniel, C. W. V. 2020. Sexually selected traits as bioindicators?: exposure to mercury affects carotenoid-based male bill color in zebra finches. - Ecotoxicology 29: 1138–1147.

Stanczyk, F. Z., Petra, P. H., Senner, J. W. and Novy, M. J. 1985. Effect of dexamethasone treatment on sex steroid-binding protein, corticosteroid-binding globulin, and steroid hormones in cycling rhesus macaques. - Am. J. Obstet. Gynecol. 151: 464–470.

Stojanovic, D., Webb nee Voogdt, J., Webb, M., Cook, H. and Heinsohn, R. 2016. Loss of habitat for a secondary cavity nesting bird after wildfire. - For. Ecol. Manage. 360: 235–241.

Trenberth, K. E., Dai, A., Van Der Schrier, G., Jones, P. D., Barichivich, J., Briffa, K. R. and Sheffield, J. 2014. Global warming and changes in drought. - Nat. Clim. Chang. 4: 17–22.

Varian-Ramos, C. W., Karubian, J., Talbott, V., Tapia, I. and Webster, M. S. 2010. Passerine, Offspring sex ratios reflect lack of repayment by auxiliary males in a cooperatively breeding. - Behav. Ecol. Sociobiol. 64: 967–977.

Vernasco, B. J., Horton, B. M., Ryder, T. B. and Moore, I. T. 2019. Sampling baseline androgens in free-living passerines: Methodological considerations and solutions. - Gen. Comp. Endocrinol. 273: 202–208.

Webster, M. S., Varian, C. W. and Karubian, J. 2008. Plumage color and reproduction in the red-backed fairy-wren: Why be a dull breeder? - Behav. Ecol. 19: 517–524.

Wingfield, J. C. 2013. The comparative biology of environmental stress?: behavioural endocrinology and variation in ability to cope with novel, changing environments. - Anim. Behav. 85: 1127–1133.

Wingfield, J. C. and Kitaysky, A. S. 2002. Endocrine responses to unpredictable environmental events: Stress or anti-stress hormones? - Integr. Comp. Biol. 42: 600–609.

Wingfield, J. C., Maney, D. L., Breuner, C. W., Jacobs, J. D., Lynn, S., Ramenofsky, M. and Richardson, R. D. 1998. Ecological bases of hormone-behavior interactions: The emergency life history stage. - Am. Zool. 38: 191–206.

Wingfield, J. C., Lynn, S. E. and Soma, K. K. 2001. Avoiding the “costs” of testosterone: Ecological bases of hormone-behavior interactions. - Brain. Behav. Evol. 57: 239– 251.

Wingfield, J. C., Pérez, J. H., Krause, J. S., Word, K. R., González-Gómez, P. L., Lisovski, S. and Chmura, H. E. 2017a. How birds cope physiologically and behaviourally with extreme climatic events. - Philos. Trans. R. Soc. B Biol. Sci. 372: 20160140.

Wingfield, J. C., Pérez, J. H., Krause, J. S., Word, K. R., González-Gómez, P. L., Lisovski, S. and Chmura, H. E. 2017b. How birds cope physiologically and behaviourally with extreme climatic events. - Philos. Trans. R. Soc. B Biol. Sci. 372: 20160140.

Woinarski, J. C. Z. 1999. Fire and Australian birds: a review. - Aust. Biodivers. responses to fire. Plants, birds Invertebr.: 55–112.

Woinarski, J. C. Z. and Legge, S. 2013. The impacts of fire on birds in Australia’s tropical savannas. - Emu 113: 319–352.

Zuk, M. and Kolluru, G. 1998. Exploitation of sexual signals by predators and parasitoids. - Q. Rev. Biol. 73: 415–438.

